# Genetic code degeneracy is established by the decoding center of the ribosome

**DOI:** 10.1101/2021.05.16.444340

**Authors:** Shixin Ye, Jean Lehmann

## Abstract

The degeneracy of the genetic code confers a wide array of properties to coding sequences. Yet, its origin is still unclear. A structural analysis has shown that the stability of the Watson-Crick base pair at the second position of the anticodon-codon interaction is a critical parameter controlling the extent of non-specific pairings accepted at the third position by the ribosome, a flexibility at the root of degeneracy. Based on recent cryo-EM analyses, the present work shows that residue A1493 of the decoding center provides a significant contribution to the stability of this base pair, revealing that the ribosome is directly involved in the establishment of degeneracy. Building on existing evolutionary models, we show the evidence that the early appearance of A1493 and A1492 established the basis of degeneracy when an elementary kinetic scheme of translation was prevailing. Logical considerations on the expansion of this kinetic scheme indicate that the acquisition of the peptidyl transferase center was the next major evolutionary step, while the induced-fit mechanism, that enables a sharp selection of the tRNAs, necessarily arose later when G530 was acquired by the decoding center.

## INTRODUCTION

Two types of degeneracy families are essentially present in the genetic code table: fourfold degenerate families and two-fold degenerate families. Degeneracy stems from the tolerance of non-Watson-Crick (WC) base pairs at the third position of the codons. In mitochondria and other small genome entities, the extent of this tolerance often fully overlaps with degeneracy, implying that the number of different tRNAs required to translate all amino-acid encoding codons is minimal. Thus, in yeast and human mitochondria, all codons in any four-fold degenerate codon family are translated by a single tRNA (that most often has an unmodified U in pos. 34), while two tRNAs are required for the translation of either purine-ending or pyrimidine-ending codons in contiguous two-fold degenerate families (Bonitz et al. 1980, Suzuki et al. 2020). These two possibilities are respectively referred to as ‘superwobbling’ and ‘wobbling’ (Rogalski et al. 2008). Based on a structural analysis of parameters identified by U. Lagerkvist (Lagerkvist 1978), it was demonstrated that the level of stability of the WC geometry of the base pair at the second position of the anticodon (N_35_-N_2_) determines the distribution of these two categories of degeneracy in the entire genetic code table (Lehmann and Libchaber 2008). Three sets of hydrogen bonds contribute to the stabilization of N_35_-N_2_:

1. The number of hydrogen bonds established by the base pair itself (N_35_-N_2_), necessarily WC.
2. The number of hydrogen bonds established by the WC base pair at the first codon position (N_36_-N_1_).
3. The strong hydrogen bond between U_33_ 2’OH and N_35_, that only occurs when N_35_ is a purine (R) (Auffinger and Westhof 2001).

Considering the sum **S** of hydrogen bonds defined in 1-3, it was shown that when **S** ≤ 5, the considered codon belongs to a two-fold degenerate family, while it belongs to a four-fold degenerate family if **S** > 5 (Lehmann and Libchaber 2008). The WC geometry of N_35_-N_2_ is critical: it enables the decoding center to adopt a configuration leading to ribosome closure (Ogle et al. 2001, 2002), which triggers GTP hydrolysis on EF-Tu and the subsequent release of the tRNA for accommodation (Voorhees et al. 2010). This geometry can be perturbed by non-WC base pairs at the third position of the codons. The model shows that penalizing N_34_-N_3_ mismatches can sufficiently alter that geometry to prevent the decoding center from adopting a productive configuration. With **S** > 5, any perturbation by the four possible base pairs at the third position is contained by N_35_-N_2_, and superwobbling is possible, whereas base pairing is restricted to simple wobbling when **S** ≤ 5, which has allowed the encoding of two different amino acids (or an amino acid and the stop function) by the considered N_1_N_2_ doublet during the expansion of the initial genetic code. At the time when this model was published, the dynamics of the decoding center was unknown, and its three residues (A1493, A1492 and G530) were assumed to be either all in the OFF or all in the ON state (resp. *syn* and *anti* for G530), the latter case corresponding to a situation where they are tightly packed and form hydrogen bonds along the minor groove of the anticodon-codon complex. In that state, the ribosome is engaged to accept the tRNA (Ogle et al. 2001, 2002, Schmeing et al. 2009, Voorhees et al. 2010). This *a priori* type of dynamics implied that an essential aspect of the model was unsatisfactory: in the all-OFF state, the respective contributions of the hydrogen bonds of N_36_-N_1_ and N_35_-N_2_ to the stability of the N_35_-N_2_ base pair were identical, which was physically implausible (a remarkable property of the parameters is that only their sum determines degeneracy, implying that they are *equivalent*). To resolve this inconsistency, it was envisioned (although not clearly stated) that residue A1493 would *always* bind to the minor groove of N_36_-N_1_ when N_36_-N_1_ and N_35_-N_2_ were complementary, even in the occurrence of penalizing mismatch at the third position. This binding (A minor, type I) is stronger with G_36_-C_1_ or C_36_-G_1_ as compared to A_36_-U_1_ or U_36_-A_1_, thereby amplifying the difference already present between these pairs. A structural context with N_36_-N_1_ as a triple base pair (N_36_-N_1_-A_1493_) would explain why N_35_-N_2_ and N_36_-N_1_ had an apparent similar weight in the stability of the N_35_-N_2_ base pair. It implied, however, that the decoding center would be already partially ON even though the tRNA could still be rejected by the ribosome.

Here we show that the possibility of the ‘partially ON’ configuration of the decoding center is confirmed by cryo-EM analyses of Loveland et al. (2017) and Fislage et al. (2018), which allows us to strengthen and extend the conclusions of the initial analysis (Lehmann and Libchaber 2008). These studies identified three different stages of the decoding center in the timeline from initial tRNA binding down to ribosome closure. In the intermediate stage, during which a tRNA is tested for anticodon-codon complementarity by the decoding center, all three examined configurations (cognate and near-cognate with either G_35_-U_2_ or A_36_-C_1_ mismatch) show that A1493 is in minor groove binding position, with a clear binding occurring in the cognate and G_35_-U_2_ cases. These new data allow us to reanalyse the specific roles of all three residues of the decoding center in terms of their contributions to both degeneracy and induced fit. In agreement with evolutionary models, we show that their dynamics suggests an early appearance of A1493 and A1492 on the ribosome at a time when no catalytic site was present and when an early kinetic scheme of translation that did not include tRNA accommodation was prevailing. In this early kinetic scheme, inferred from a physico-chemical correlation in the genetic code, our analysis suggests that the initial role of A1493 and A1492 was to allow a relaxation of base pairing specificity at the third position of the codons through the compensatory strengthening they implemented at the first position, which gave rise to degeneracy. Kinetics considerations suggest that the peptidyl transferase center (PTC) was the next major acquisition by the ribosome, while proofreading (Hopfield 1974, Ninio 1975, Thompson and Stone 1977) arose at a later stage with the initial form of EF-Tu•GTP. It logically follows that the controlled hydrolysis of EF-Tu’s GTP through 30S closure by induced fit was a latecomer mechanism, implemented when G530 was acquired by the decoding center.

## RESULTS

### The structural model of degeneracy is consistent with cryo-EM data

Recent cryo-EM investigations on the decoding mechanism of the ribosome have allowed the identification of three different states of the decoding center and the A-site tRNA in the timeline from initial tRNA ribosome binding down to 30S closure (Loveland et al. 2017, Fislage et al. 2018). Following Fislage et al.’s notations (see Figure 7 of their publication), these states are: initial tRNA binding, tRNA sampling and engaged state, the latter state corresponding to a closed 30S subunit, in which the ribosome commits to accept a tRNA. The structures show that with a single mismatch at either the first or the second position of the codon, or in the cognate case, residue A1493 moves to and remains in the ‘ON’ position during tRNA sampling, i.e. flipped out of helix 44 and in N_36_-N_1_ minor groove binding position. With a A_36_-C_1_ mismatch at the first position, A1493 does not form hydrogen bonds with the minor groove, no AC pair being formed (Fig. 1A). With a G_35_-U_2_ mismatch at the second position, a A_36_-U_1_ base pair does form, and A1493 binds to its minor groove, although none of its three hydrogen bonds is optimal (Fig. 1B). In the cognate case, A1493 binds to the minor groove and forms h-bonds during tRNA sampling (Fig. 1C). There is no existing structure with a forbidden base pair at the third position only, for which the model predicts that A1493 would, likewise, bind to the first base pair during tRNA sampling. The above data, however, clearly support this possibility.

**Figure 1.**
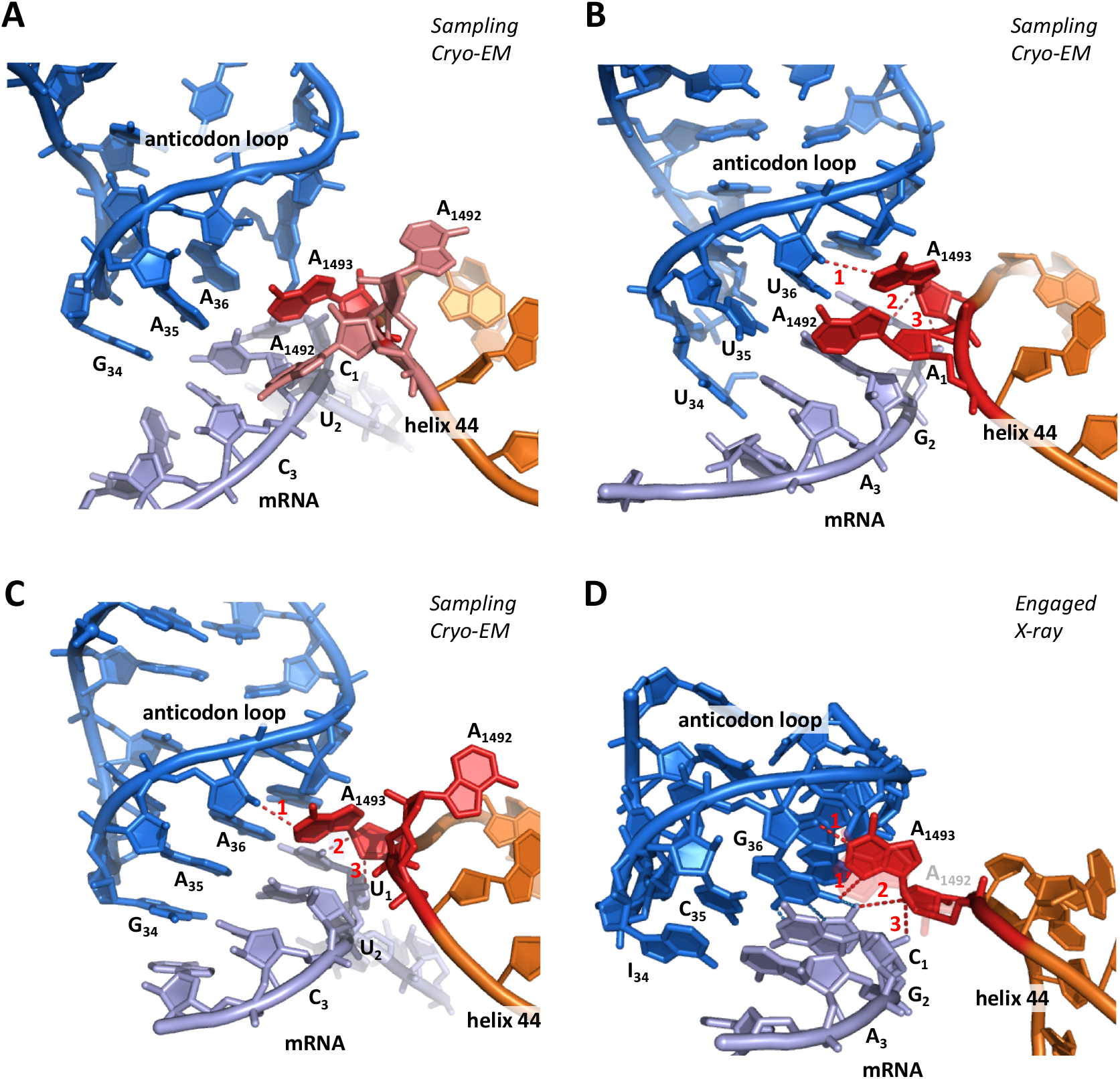
**Cryo-EM (A-C) and X-ray (D) structures of anticodon-mRNA complex within the decoding center of the ribosome (for clarity, G530 and helix h18 are not shown). A) Non-cognate interaction, with AC mismatch at the first position in the state of tRNA sampling (pdb 5wfk, Fislage et al. 2018). Although A1493 is ON, no hydrogen bond with the minor groove can occur. CryoEM resolution is 3**.**4 Å. B) Non-cognate interaction, with GU mismatch at the second position in the state of tRNA sampling (pdb 5uyp, Loveland et al. 2017). A1493 binds to the minor groove. Hydrogen bond D-A lengths are 1: 3**.**6 Å; 2: 3**.**0 Å; 3: 4**.**5 Å (avg**.**: 3**.**7 Å). CryoEM resolution is 3**.**9 Å. C) Cognate interaction in the state of tRNA sampling (pdb 5uyl, Loveland et al. 2017). A1493 binds to the minor groove. Hydrogen bond D-A lengths are 1: 3**.**0 Å; 2: 3**.**1 Å; 3: 3**.**8 Å (avg**.**: 3**.**3 Å). CryoEM resolution is 3**.**6 Å. D) X-ray structure of a cognate interaction (pdb 1xnq, Murphy and Ramakrishnan 2004) illustrating an A minor interaction with a GC base pair at the first position. Hydrogen bond D-A lengths are 1: 2**.**6 Å; 1’: 2**.**9 Å; 2: 3**.**3 Å; 3: 2**.**5 Å (avg**.**: 2**.**8 Å). Compared to pdb 5uyl, examination of the 5uym pdb structure suggests that the shorter length of these bonds results from A1493 and A1492 being both bound to the anticodon-codon complex. Xray resolution is 3**.**05 Å. In order to highlight hydrogen bonds, the angle of view was tilted compared to the other structures, and A1492 is semi-transparent. Overall, A1492 is found about 50% of the time in the ‘ON’ state during tRNA sampling (Fislage et al. 2018). Specific densities of A1492 are such that it is 50% ON/50% OFF in the 5wfk structure (light pink), ON in the 5uyp structure (red) and OFF in the 5uyl structure (red)**.

Because the wobble position is two base pairs away from the A1493 binding site, a N_34_-N_3_ mismatch generates a smaller perturbation at the A1493 binding site than a N_35_-N_2_ mismatch, for which A1493 A minor binding during tRNA sampling is now confirmed (Fig. 1B). A complete demonstration would, however, require a structure with a base pair more penalizing than G_34_-U_3_ at the third position, e.g. U_34_-U_3_ or U_34_-C_3_ (U_34_ is almost always involved in superwobbling; Bonitz et al. 1980). In brief, cryo-EM analyses have revealed that the A1493 residue of the decoding center binds to the minor groove of N_36_-N_1_ during tRNA sampling if this base pair is Watson-Crick, a binding that *further stabilizes the complex* during the time it is tested by residues A1492 and G530 for 30S closure.

### Degeneracy in the genetic code is established through a major contribution by A1493

The cryo-EM data of Loveland et al. and Fislage et al. allow us to refine the structural model of degeneracy previously described (Lehmann and Libchaber 2008). Figure 2A highlights the four different levels specifying the stability of the WC geometry of the N_35_-N_2_ base pair during tRNA sampling in the situation when both N_36_-N_1_ and N_35_-N_2_ are complementary. The two lowest levels attribute a two-fold degeneracy to the corresponding codons, while the two highest levels attribute a four-fold degeneracy. As a result of the equivalence of Lagerkvist’s parameters, levels 2 and 3 are degenerate in such a way that three configurations of hydrogen bonding patterns are possible. Remarkably, to each configuration correspond two sets of codons related by A_1_ ↔ U_1_ or G_1_ ↔ C_1_ permutations (indicated on each anticodon stem in Fig. 2A). Consequently, when A_1_ (G_1_) and U_1_ (C_1_) are mirror ordered with respect to the center of the table (dashed line), the two degeneracy families are also symmetrically arranged with respect to the center (Fig. 2B).

**Figure 2.**
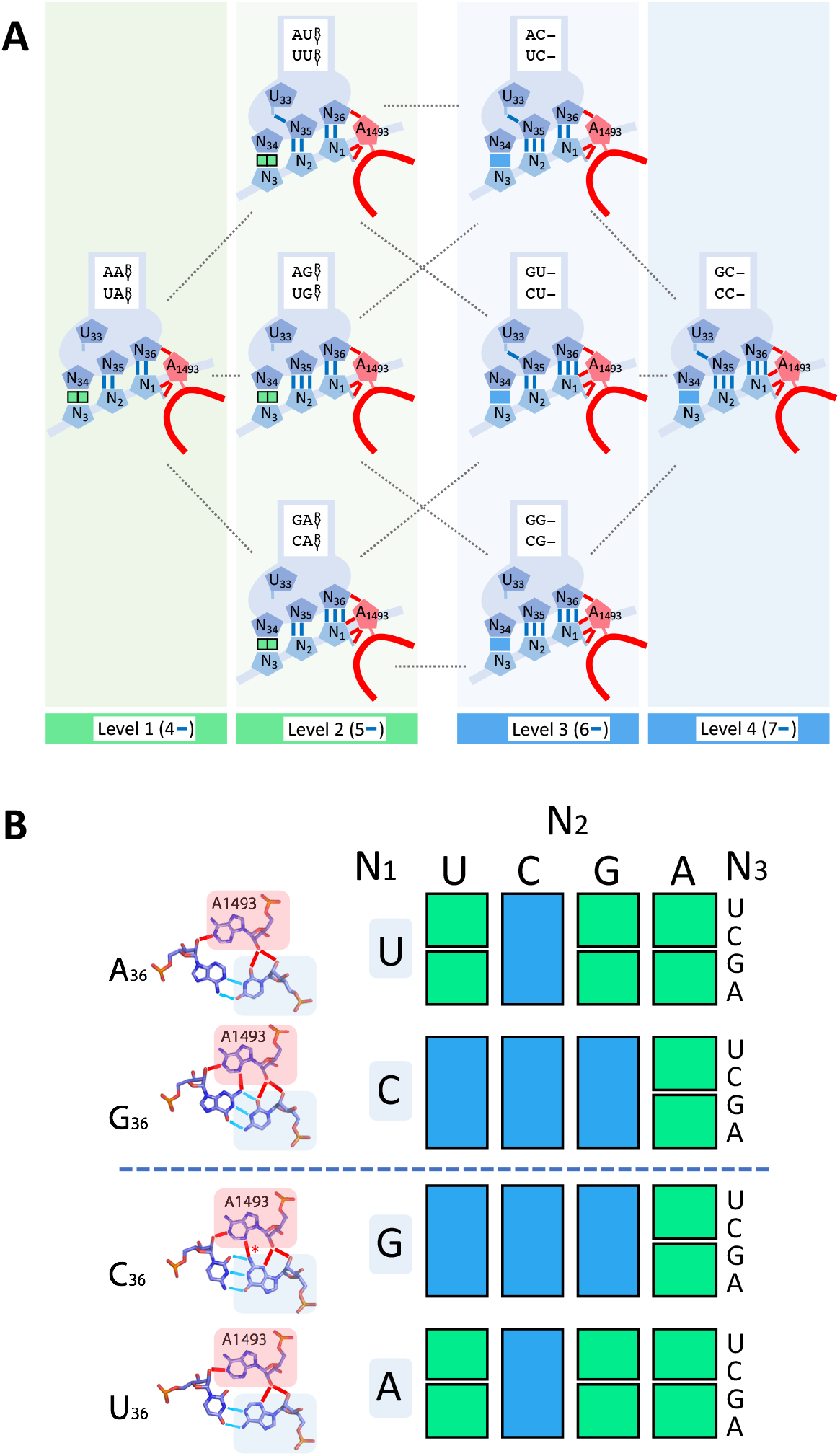
**Relation between hydrogen bonding patterns involved in the stability of the WC geometry of N**_**35**_**-N**_**2**_ **and degeneracy. A) Levels of stability of the WC geometry of the N**_**35**_**-N**_**2**_ **base pair during tRNA sampling, as determined by hydrogen bonds associated with Lagerkvist’s parameters (in blue) and residue A1493 (in red). Levels 1 and 2 specify two-fold degenerate families (contiguous green boxes), while levels 3 and 4 specify four-fold degenerate families (blue boxes). B) Yeast or human mitochondria genetic code table highlighting the two families of degeneracy (same color code as in A). Amino acids are not specified to point out that they are not primarily involved in the determination of these families. The A minor interaction between A1493 and N**_**36**_**-N**_**1**_ **is shown on the left. All shown hydrogen bonding patterns were found in experimental structures (see Fig. 1), except that of C**_**36**_**-G**_**1**_**-A**_**1493**_, **for which no structure could be identified in the pdb database. In that case, the only hypothetical hydrogen bond, highlighted with an asterisk*, is expected to occur similarly as for the G**_**36**_**-C**_**1**_**-A**_**1493**_ **configuration due to the position of the G**_**1/36**_**(C**_**1**_**-NH**_**2**_**) amino group at the center of the base pair**.

According to the analysis, the most remarkable effect that occurs when both N_36_-N_1_ and N_35_-N_2_ are complementary is the *positive* selection of tRNAs enforced by A1493: the strengthening of N_35_-N_2_ resulting from N_36_-N_1_ A minor binding enables the acceptance of some tRNAs with non-WC base pairs at the third position, whereas tRNAs are counterselected when N_36_-N_1_ and/or N_35_-N_2_ are not complementary (Ogle et al. 2001, 2002, Loveland et al. 2017, Fislage et al. 2018). The involvement of A1493 in degeneracy provides an explanation for why Lagerkvist’s parameters are equivalents (Fig. 2A): each increase in the level of stability of N_35_-N_2_ occurs upon the addition of either 1 *local* hydrogen bond (U_33_-N_35_ or N_35_-N_2_, in blue) or 2 hydrogen bonds on the *neighboring* triple base pair (N_36_-N_1_-A_1493_, one in blue and one in red), revealing that these two possibilities are equivalent in term of the added stability to N_35_-N_2_.

A striking aspect of the model is that no stacking parameter is required. It suggests that the high number of hydrogen bonds involved (7 to 11) confer structural energies that dominate over the variability of the stacking interaction, which further corroborates the implication of A1493 in degeneracy. The number of these hydrogen bonds is invariant upon A_1_ ↔ U_1_ or G_1_ ↔ C_1_ permutations. In the case of G_36_-C_1_ and C_36_-G_1_, this property stems from the position of the G_1/36_(C_1_-NH_2_) amino group at the center of the base pair (Figs. 1D and 2B). Although stacking is not a parameter, N_37_ stabilizes the N_36_-N_1_ base pair by stacking on it, an effect that is optimal since this base is a conserved purine (Auffinger and Westhof 2001). Stabilization is further enhanced when N_37_ is modified (Grosjean et al. 1998; Konevega et al. 2004, Jenner et al. 2010, Grosjean and Westhof 2016), and the extent of modification negatively correlates with the G+C composition of the anticodon (Grosjean et al. 1998, Grosjean and Westhof 2016), indicating that this base also contributes to an adjustment of the overall stability of each anticodon-codon interaction, and is thus likely a hidden requirement to the observed degeneracy. Deformation of the tRNA body has also been shown to affect the extent of wobbling at the third position (see summary and discussion Section). With regard to the present analysis, the *directional* nature of hydrogen bonds plausibly explains why they play a predominant role in the stability of the *geometry* of the N_35_-N_2_ WC base pair, which is the decisive criteria for ribosome closure (Loveland et al. 2017, Fislage et al. 2018). In a situation relevant to degeneracy (Fig. 2A), this geometry is preserved if the network of hydrogen bonds stabilizing N_35_-N_2_ is strong enough to contain the perturbation generated by a given non-canonical N_34_-N_3_ base pair.

### The induced-fit mechanism is a late acquisition of the decoding center

The implication of A1493 in degeneracy shows that the implementation of unspecific pairing at the third position of the codons arose at the time when the ribosome acquired residue A1493 on helix h44. Remarkably, two analyses suggest that the segment of h44 where A1493 and A1492 are located appeared early in the evolution of the ribosome, whereas helix h18, harboring G530, emerged at a much later stage (Harish and Caetano-Anollés 2012, Petrov et al. 2015) (Fig. 3A). The latter residue has a major role in the induced-fit mechanism: it drives 30S closure (Loveland et al. 2017, Fislage et al. 2018), which triggers GTP hydrolysis on EF-Tu, thereby releasing the incoming tRNA for accommodation (Voorhees et al. 2010, Kavaliauskas et al. 2018). The mentioned models on ribosome evolution are thus consistent with the induced fit of the decoding center being, logically, established later than degeneracy. The connection between the successive emergence of helices h44 and h18 and these fundamental aspects of translation must be underscored. The mechanism itself reflects this evolutionary succession:

**Figure 3.**
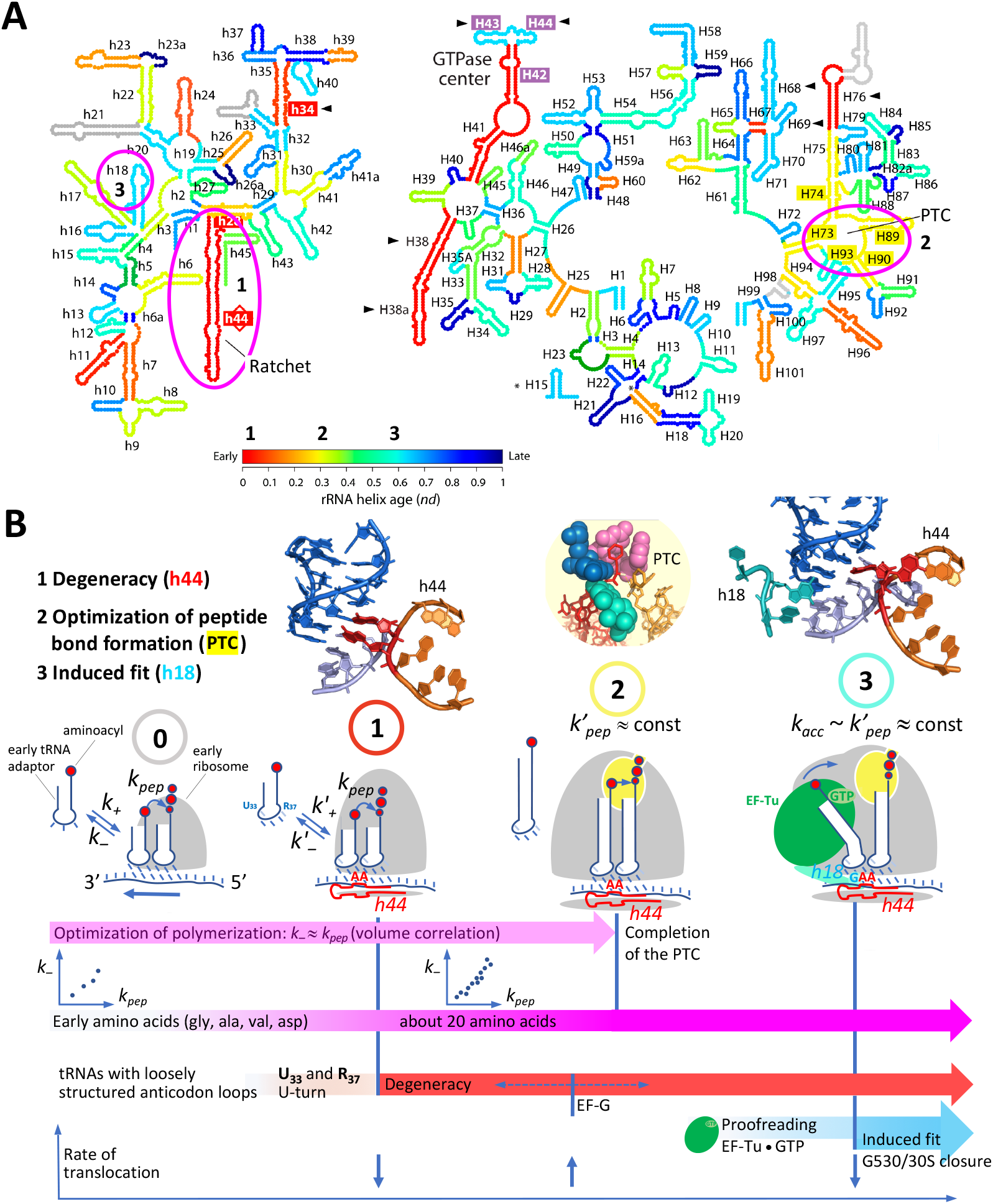
**Evolution of rRNA structures in the model of Harish and Caetano-Anollés and evolution of decoding in translation based on the analysis of degeneracy. A) rRNA evolution. Three specific helices (or groups of helices) involved in transitions in the evolutionary model of decoding are highlighted. Adapted from Harish and Caetano-Anollés (2012). B) Evolutionary model of decoding on the ribosome. From the origin until the advent of the PTC, a Michaelis-Menten type of kinetic inferred from the volume correlation (Lehmann 2000) governs the rate of translation, with tRNA association (*k***_***+***_**) and dissociation (*k***_***-***_**) rate constants, and a kinetic constant of peptide bond formation (*k***_***pep***_**), sometimes called *k***_***cat***_ **in earliers works (Lehmann 2000, 2018, Lehmann et al. 2009). The advent of U**_**33**_ **and R**_**37**_, **as well as helix h44 (A1493 & A1492) modulated these kinetic constants (*k*′**_***+***_, ***k*′**_***-***_**). Relevant structural contexts are shown above each evolutionary transition: decoding center with h44 only (1), peptidyl transferase center (PTC) (2) and whole decoding center with helix h18 (3). See text for additional explanations. Note that all three considered transitions highlighted in A and B concur, although these two models were established essentially independently**.

A1493 *first* binds to the minor groove, while A1492 fluctuates between ON and OFF states; *only then* can A1492 and G530 fully bind to the complex in the cognate case, thereby achieving ribosome closure (Loveland et al. 2017, Fislage et al. 2018).

### The appearance of A1493 generated a decoding transition on the ribosome

The involvement of A1493 in degeneracy highlighted by the present analysis, and the coherence of the sequential buildup of the decoding center in the evolutionary models of Harish and Caetano-Anollés (2012) and Petrov et al. (2015) motivated us to outline a model of evolution of ribosomal decoding based on the identified role of A1493 and a plausible form of the earliest kinetic scheme of translation (Lehmann et al. 2009). This kinetic scheme (Fig. 3B, left) was established from an interpretation of a physico-chemical correlation in the genetic code called the volume correlation (Lehmann 2000, 2017, 2018). This correlation suggests that at the origin of translation, the lifetime of the association between a tRNA and a complementary codon was about equal to the characteristic time required by the aminoacyl carried by this tRNA to make a peptide bond, which was side-chain dependent. This adjustment, which can be expressed with kinetic constants as ***k-*** _anticodon-codon_ ≈ ***k***_***pep*** aminoacyl_, implies that the aminoacyls were in immediate position for forming a peptide bond upon tRNA codon binding —i.e. there was no tRNA accommodation at the origin— while not being confined inside a catalytic site, which would have standardized the ***k***_***pep*** aminoacyl_s to an approximately uniform value, an action that is achieved by the peptidyl transferase center (PTC) of modern ribosomes (Lehmann 2017). An elementary Michaelis-Menten kinetic scheme comprising the above kinetic constants best encapsulates these features (Fig. 3B, left). An analysis shows that in this model, the rate of translation is optimal precisely when ***k-*** _anticodon-codon_ ≈ ***k***_***pep*** aminoacyl_ occurs for all tRNA:aminoacyl couples (Cibils et al. *in prep*.). As this analysis has not yet been published, this property is left here as a conjecture.

In the context defined by this model of the early translation, a straightforward consequence of the strengthening of the N_36_-N_1_ base pair that occurred when A1493 became functional on h44 was a relaxation of base pairing specificity at the third position of the codons, a rebalancing scheme that would have overall preserved the ***k-*** _anticodon-codon_s, and thus the rate of translation. In a context of a limited variety of tRNAs, this action of A1493 presumably led to an increase in the processivity and accuracy of translation, discussed below.

Because a mismatch perturbs the geometry and stability of neighboring base pairs along a double helix, the type I A minor binding achieved by A1493 could have been optimal only at the first position, i.e. two base pairs away from the third position (where tolerated mismatches would occur), which may explain why this solution was selected.

Although current models of ribosome evolution may not predict whether A1493 and A1492 were both initially present on h44 (Harish and Caetano-Anollés 2012, Petrov et al. 2015), this possibility is plausible since the dynamics of A1493 would likely be altered without A1492, and the type II A minor binding achieved by A1492 (Ogle et al. 2001), which is more tolerant to mismatch (it does not bridge over N_35_-N_2_), may contribute to N_36_-N_1_ stabilization. This binding occurs ∼50 % of the time during tRNA sampling (Fislage et al. 2018). In that state, the tRNA is partially bent, a feature associated with presence of EF-Tu that allows an optimal substrate selection through deformation (Yarus et al. 2003, Savir and Tlusty 2007, Schmeing et al. 2009, Schmeing et al. 2011, Savir and Tlusty 2013). Because EF-Tu, an elaborate protein cofactor, could not have occurred at the origin of translation (which is consistent with an absence of tRNA accommodation, inferred from the volume correlation), it can be maintained that the stem of initial tRNA adapters did not undergo such deformation. In that case, A1492 would bind 100% of the time to the complex upon tRNA codon association, similarly as it does with fully accommodated tRNAs on modern ribosomes. A fully bound A1492 may contribute to an optimal strengthening mediated by A1493 in the situation when both N_36_-N_1_ and N_35_-N_2_ are Watson-Crick.

Structural and functional considerations suggest that both the processivity and accuracy of translation increased when residues A1493 and A1492 became functional on helix h44:

#### Processivity of translation

In the proposed Michaelis-Menten kinetic scheme of the initial translation (Fig 3B), the relaxation in base pairing specificity that occurred at the third position of the codons through the action of A1493 and A1492 may have allowed a given set of different transfer tRNAs, necessarily limited at the origin, to be more tolerant to mutations incorporated at the third position during replication (Fig. 4), and thus translate longer sequences.

**Figure 4.**
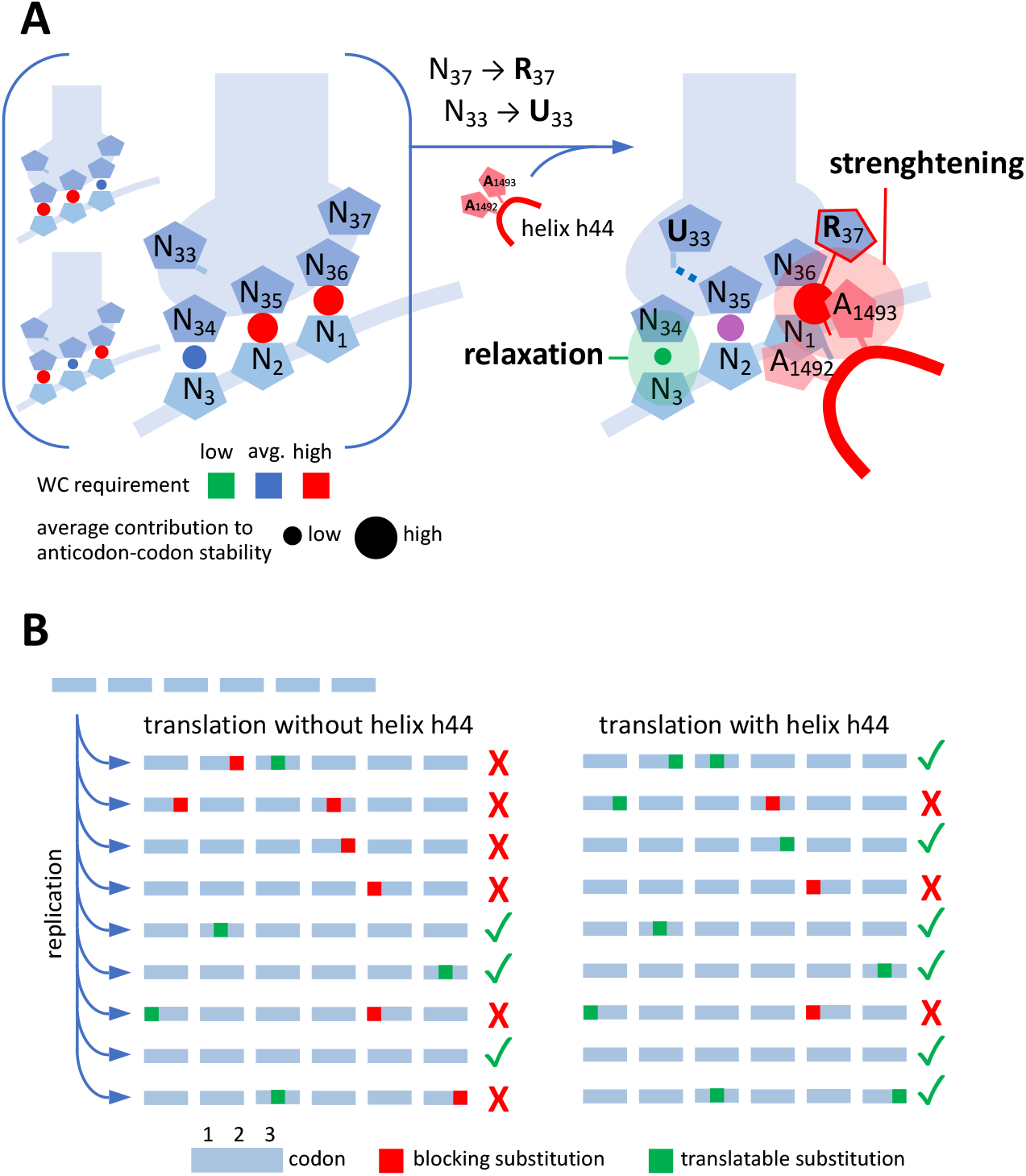
**Evolutionary transition 1: from early tRNA anticodon loops and no decoding center to U**_**33**_ **and R**_**37**_ **-shaped anticodon loops and helix h44 on the ribosome (A1493 and A1492). A) Left: initial loop of tRNA adapter, with little structuration, bound to a codon in an absence of decoding center. Although the shape of the loop might provide a high flexibility to the base pair at the 3**^**rd**^ **position, single GU wobble base pairs could occur in pos. 2 or 1 (background) while still providing enough stability to ensure peptide bond formation in the early translation mechanism. Right: the advent of R**_**37**_ **and helix h44 strengthened the anticodon-codon interaction at the 1**^**st**^ **position, while the U-turn (U**_**33**_**) helped relax base pairing specificity at the 3**^**rd**^ **position. R**_**37**_ **stacking on N**_**36**_**-N**_**1**_ **is schematized with a thin red line. B) Translation of early coding sequences: suggested improved processivity resulting from transition 1. Because the early replication mechanism is inaccurate, RNA sequences accumulate mutations, and thus may not always be fully translated due to reduced sets of tRNAs (left). The advent of h44 together with anticodon loop structuration (see A) provided an improved processivity during translation by lowering base pairing requirement at the third position (right)**.

As there was initially no strong geometrical requirement for the base pair at the second position in the absence of G530 and induced-fit mechanism, unspecific base pairing at the third position, that perturb the N_35_-N_2_ geometry, was plausibly less stringent than that occurring on modern ribosomes. A1493 and A1492 binding would compensate for the loss in anticodon-codon stability generated by mismatches at the third position within a simple rebalancing scheme (Fig. 4A).

It suggests to us that the four codons belonging to any of the 16 N_1_N_2_ doublets of the genetic code may have been translated by a single tRNA upon the action of h44 in an all four-fold-degeneracy regime following transition 1 (Fig. 5, center). This possibility does naturally not imply that all 16 doublets were encoding amino acids, at least immediately following this early transition. It is striking that the acquisition of R_37_ and U_33_ on the anticodon loop, that presumably also occurred early in the evolution of the tRNAs, respectively reinforced the N_36_-N_1_ base pair through R_37_ stacking and provided an extended conformational freedom to N_34_ at the edge of the U-turn (Quigley and Rich 1976), in an apparent synergism with the effect of h44 (Fig. 4A). The early replication mechanism being inaccurate, the arising degeneracy most likely improved the processivity of translation among mutated copies of early RNA genes (Fig. 4B). In the absence of decoding center, wobbling could occur at any codon position at the origin of translation (Fig. 4A, left), and thus lead to a miscoding that would be prohibitive to the emergence of Life. It has been suggested that a very limited codon and anticodon repertoire such as the ‘GNC’ code (where N is A, G, C or U) could overcome this issue while simultaneously managing frameshifting and frame indeterminacy at that stage (Eigen and Schuster 1978, Ikehara et al. 2002, Wang and Lehmann 2016) (Fig. 5, left).

**Figure 5.**
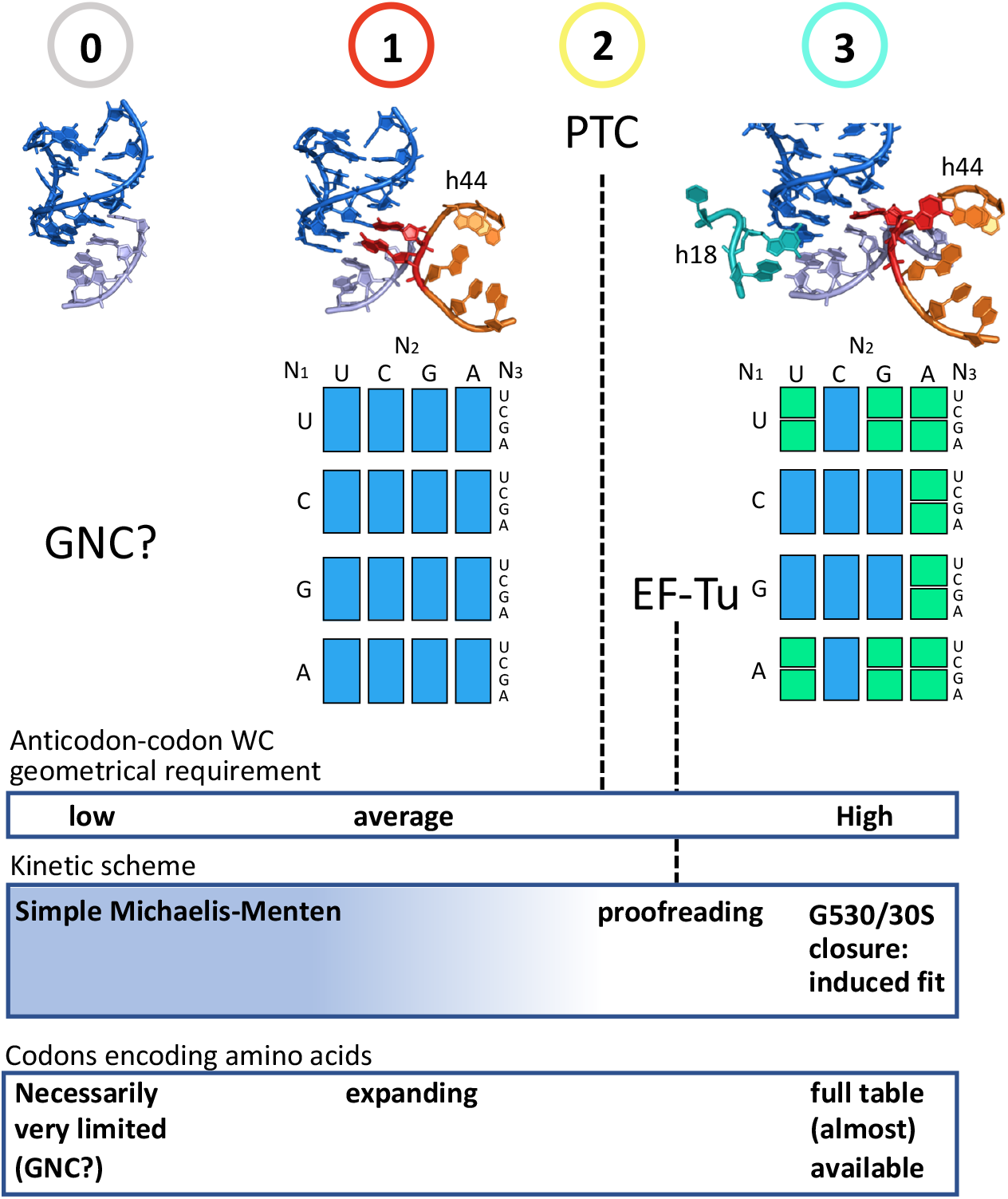
**Anticodon-codon interaction and codon degeneracy in the genetic code during ribosome evolution. From an early hypothetical structure with no decoding center (initial state, 0), in which the properties of a GNC code may have provided a required stability to an early translation system (Eigen and Schuster 1978, Wang and Lehmann 2016), evolutionary models and the dynamics of the decoding center suggest that helix h44 with A1493 and A1492 appeared first (transition 1), which enabled an extended degeneracy at the third position (blue boxes). The completion of the PTC (transition 2) and the appearance of EF-Tu (proofreading) necessarily occurred before a controlled hydrolysis on EF-Tu by the decoding center through G530 and 30S closure (transition 3), which gave rise to modern degeneracy. Inferred kinetic scheme and codons occurring from stage 0 to transition 3 are indicated at the bottom**.

#### Accuracy of translation

a GU wobble base pair is only slightly less stable than an AU base pair (Freier et al. 1986), but it is *tilted* compared to a regular WC base pair. In the context of the N_36_-N_1_-A_1493_ triple base pair, no such degree of freedom is available due to *planar* constraints: in order for A1493 to establish optimal hydrogen bonds with N_36_-N_1_, this base pair has to display a WC geometry (Ogle et al. 2001, Ogle et al. 2002). Thus, an increase in the dimensionality of the anticodon-codon complex is associated with an increased selectivity. Furthermore, Satpati et al. (2014) found out through molecular dynamic simulations that mismatches are penalized essentially as a result of water exclusion due to the binding of A1493, A1492 and G530: missing hydrogen bonds occurring in mismatches cannot be compensated through hydrogen bonding with water. This effect could already partially occur without loop h18 and G530.

Another effect resulting from the action of h44 must be considered: because the A-site tRNA and the RNA template became caught by A1493 and A1492 upon anticodon binding, the rate of translocation necessarily slowed down (Fig. 3B, bottom). On modern ribosomes, the grip of the decoding center constitutes a barrier to translocation, which is overcome by the elongation factor EF-G and the free-energy available from the hydrolysis of a GTP (Katunin et al. 2002, Frank et al. 2007; Taylor et al. 2007, Liu et al. 2014). Without helix h18 and G530, translocation could still spontaneously occur through thermal fluctuations (Ling and Ermolenko 2016). A consistent evolutionary scenario is that an ancestor of EF-G came into the picture after the emergence of A1493 and A1492, which would alleviate the early grip, and make the second transition to G530 and proofreading possible by preventing a catastrophic slowdown of translocation upon building of the full decoding center (Fig. 3B, bottom). During evolution, an early fixation of R_37_, which makes an interstrand stacking and thus helps maintain the reading frame, would also best ensure the maintenance of that frame upon appearance of A1493 and A1492 (Figs. 3B and 4A). The subsequent appearance of EF-G and R_37_ modifications would further reduce frameshifting events during translocation (Konevega et al. 2004, Jenner et al. 2010, Liu et al. 2014, Zhou et al. 2019, Peng et al. 2019).

### Co-evolution of the translation machinery and the genetic code

This section summarizes and brings further justifications to the evolutionary model depicted in Figure 3B. Remarkably, all three major transitions highlighted in this scenario agree with the model of ribosome evolution proposed by Harish and Caetano-Anollés (2012) (Fig. 3A). While still being consistent with the model of Petrov et al. (2015), our analysis does not support a very early appearance of the PTC on the ribosome, as suggested by this study (see summary and discussion Section).

#### Initial stage (0)

although no strong evidence so far explains the origin of RNA and how the initial translation came about, the volume correlation in the genetic code (Lehmann 2000) suggests that the early translation was driven by a simple Michaelis-Menten kinetic scheme (Lehmann et al. 2009, Lehmann 2017, 2018). The fixation of U_33_ and R_37_ on the anticodon loops, that improved anticodon-codon associations and helped maintain the reading frame (Konevega et al. 2004), was plausibly an early acquisition on all tRNAs.

#### First major transition (1)

residues A1493 and A1492 appeared on helix h44. Together with U_33_ and R_37_, they established the basis of modern degeneracy (Fig. 4 and Fig. 5, center).

#### Second major transition (2)

build-up of the PTC. Because this catalytic site confines the aminoacyls in a desolvated environment, the amino groups are more reactive (Johansson et al. 2011). Furthermore, an induced-fit mechanism orients the aminoacyls for nucleophilic attack, which cancels the conformational freedom available to the amino group in solution, that is side-chain dependent (Lehmann 2017). As a consequence, all ***k***_***pep*** aminoacyl_s are levelled up to an approximately uniform ***k’***_***pep***_ value. Thus, at the time of the completion of the PTC, the [***k-*** _anticodon-codon_ ≈ ***k***_***pep*** aminoacyl_] optimization that had guided the establishment of the code became obsolete. Free from this constraint, the genetic code could evolve on its own, although codon reassignment is known to have occurred at an extremely low rate –otherwise, the volume correlation would have disappeared.

Because it would break the initial simple MM kinetic scheme (Fig. 3B, left), the EF-Tu cofactor could come into the picture only after the optimization of the ***k***_***pep***_***’*** _aminoacyl_ achieved by the PTC. In the absence of G530 and an induced-fit mechanism, an elementary form of proofreading would occur: most plausibly, GTP hydrolysis on EF-Tu, that leads to the release of the tRNA for accommodation (Kavaliauskas et al. 2018), was initially triggered by the docking of the tRNA•EF-Tu•GTP ternary complex onto the ribosome, following the simple clockwork mechanism envisioned by Ninio and Hopfield (Ninio 1975, Hopfield 1974, Thompson and Stone 1977), which is independent of the decoding center.

#### Third major transition (3)

appearance of helix h18 and the associated induced-fit mechanism (Pape et al. 1999), that involves G530 anticodon-codon latching and ribosome closure (Ogle et al. 2001, 2002, Voorhees et al. 2010, Loveland et al. 2017, Fislage et al. 2018). This large-scale rearrangement docks EF-Tu on the saricin loop, which triggers GTP hydrolysis (Voorhees et al. 2010, Loveland et al. 2017). From the early simple proofreading mechanism (see above), a plausible evolutionary transition was a change in the structure of EF-Tu that set GTP hydrolysis under the conditional control of ribosome closure, thus *combining* induced fit with proofreading. Available data (Johansson et al. 2011, Juette et al. 2016) suggest that the kinetic constant of accommodation (***k***_***acc***_) is of the same order of magnitude as ***k’***_***pep***_ at physiological pH on modern ribosomes (Fig. 3B, right), although this point still needs to be established experimentally. Because of its sensitivity, that is tuned by tRNA deformation (Yarus et al. 2003, Schmeing et al. 2009), the induced fit would allow a much sharper discrimination between cognate and near-cognate tRNA through optimal decoding (Yarus et al. 2003, Savir and Tlusty 2007, 2013, Schmeing et al. 2011), thus giving rise to modern degeneracy (Fig. 5 right). Base modifications, that could only occur at a late stage with modifying enzymes, will still be required to shape some tRNA anticodon loops so that they can be accepted by the decoding center (Blanchet et al. 2018), best prevent leaking wobbling between contiguous 2x degenerate codon families, and ensure reading frame maintenance during translocation.

## SUMMARY AND DISCUSSION

Recent cryo-EM structures have revealed the dynamics of the decoding center of the ribosome during tRNA selection (Loveland et al. 2017, Fislage et al. 2018). Based on these results, the present work shows that residue A1493 of the decoding center plays a key role in degeneracy by strenghtening the N_36_-N_1_ base pair during tRNA sampling, which allows non-specific N_34_-N_3_ pairings to be accepted by the ribosome. This possibility was suspected at the time of an earlier work on degeneracy (Lehmann and Libchaber 2008), although it remained unclear because the dynamics of the decoding center was unknown. We now conclude that degeneracy in the modern genetic code is established by a complex comprising the anticodon, the codon and A1493, while a clear-cut distinction between contiguous two-fold degenerate families requires the induced fit mediated by the whole decoding center and modifications on the tRNA anticodon loop.

It must be emphasized that degeneracy corresponds to a maximization of wobbling (Lehmann and Libchaber 2008), which requires specific tRNAs. Decoding in mitochondria suggests that a uridine in pos. 34 can almost always achieve superwobbling in four-fold degenerate families (Bonitz et al. 1980, Rogalski et al. 2008, Suzuki et al. 2020), while some uridine modifications, such as uridine 5-oxyacetic acid, are known to further enhance this property (Näsvall et al. 2004, Weixlbaumer et al. 2007). However, most bacteria and higher order organisms use more than one tRNA to translate all codons in four-fold degenerate codon families, either by modifying U_34_ in such a way as to prevent superwobbling, by avoiding U in position 34, or by structural constraints (see below). Furthermore, in twofold degenerate codon families, U_34_ modifications (e.g., xm^5^s^2^U derivatives) are always present, and are required to prevent ‘‘leaking’’ wobbling between families sharing identical nucleotides in pos. 1 and 2 (Yokoyama et al. 1985, Yokoyama and Nishimura 1995). Codon assignment was, therefore, partially ambiguous before the appearance of modifying enzymes (and still is to some extent). The advent of inosine might explain why AUR and AUY two-fold degenerate families further reorganized into AUG and AU/U,C,A codon boxes. More generally, the extent of wobbling –and, thus, degeneracy– is controlled by structural deformations required for the anticodon to achieve proper codon binding in the context specified by the ribosome and EF-Tu (Yarus et al. 2003, Savir and Tlusty 2007, Schmeing et al. 2009, Schmeing et al. 2011, Savir and Tlusty 2013), which often requires base modifications (Blanchet et al. 2018).

Although the present analysis shows that hydrogen bonds determine the extent of degeneracy in the genetic code, experiments and molecular dynamic simulations suggest that steric complementarity between the decoding center and the anticodon-codon complex is more important than hydrogen bonds in the selection of cognate tRNAs (Khade et al. 2013, Schrode et al. 2017). There is, however, no fundamental contradiction between these two results: the network of hydrogen bonds involved in degeneracy contributes to the stabilization of the WC geometry at the second position, which is critical only when a non-canonical base pair occurs at the third position. The expected effect of missing hydrogen bonds is only a reduction of the extent of wobble base pairs accepted by the ribosome: in particular, superwobbling with U_34_ that normally occur would be prohibited when specific hydrogen bonds are missing.

In the evolutionary scenario depicted in Figure 3B, the PTC emerges after helix h44 and before helix h18, in agreement with the analysis of Harish and Caetano-Anollés (2012) (Fig. 3A). This succession can be justified by the following: because the PTC levelled the kinetic constants of peptide bond formation up to similar ***k’***_***pep*** aminoacyl_ values, it cancelled the ***k-*** _anticodon-codon_ ≈ ***k***_***pep*** aminoacyl_ adjustment that had shaped the code from the origin (Lehmann 2000, 2017, 2018). This early optimization, the trace of which is the volume correlation (Lehmann 2000), could not have occurred if the PTC was already present at the origin of translation. According to the present work, the possibility of the A1493(h44)/degeneracy rebalancing is based on this optimization, implying that h44 necessarily emerged before the PTC. Our results thus do not support an early emergence of this catalytic site, as the model of Petrov et al. (2015) suggests. Another justification of the proposed evolutionary scheme relates to tRNA accommodation, which is part of the proofreading mechanism, and implies the presence of the PTC: the tRNA acceptor arm is funnelled by rRNA helices H89 and H90-92, that are both rooted on this catalytic site (Burakovsky et al. 2010, Rakauskaitė and Dinman 2011). Also, proofreading implies a commitment of the ribosome to peptide bond formation once the 3’ end of an aminoacyl-tRNA reaches the peptidyl-tRNA, which implies high ***k’***_***pep*** aminoacyl_s of similar values, thus the PTC. We conclude that in the timeline of evolution, the completion of the PTC occurred after the appearance of degeneracy (residues A1493 & A1492) and before EF-Tu/proofreading, the induced-fit mechanism (30S closure) controlled by G530 being necessarily a latecomer.

One of the most striking structural aspect of the decoding center is that its three nucleotides are distributed on two different helices far apart from each other, implying that their simultaneous appearance in the course of the early evolution of the ribosome is highly unlikely. In agreement with evolutionary models (Harish and Caetano-Anollés 2012, Petrov et al. 2015), and with the dynamics of the decoding center (Loveland et al. 2017, Fislage et al. 2018), the major conclusion of the present analysis is that degeneracy arose when residues A1493 and A1492 took their function on helix h44 at an early stage of the evolution of the ribosome.

## Acknowledgements

We wish to thank A. Korostelev for comments and suggestions on the manuscript. We are also grateful to D. Gautheret and A. Libchaber for ongoing support.

## Author contributions

JL performed the research, discussed the results and wrote the manuscript. SY provided logistic support, discussed the results and proofread the manuscript.

## Material and methods

### Analysis of ribosome structures

Crystal and cryo-EM structures of ribosomes complexed with tRNAs or tRNA fragments (Murphy FV 4th, Ramakrishnan V. 2004, Loveland et al. 2017, Fislage et al. 2018) were retrieved from the *protein databank* website and analysed with the *Pymol* software.

